# STbayes: An R package for creating, fitting and understanding Bayesian models of social transmission

**DOI:** 10.1101/2025.06.07.658152

**Authors:** Michael Chimento, William Hoppitt

## Abstract

1. A critical consequence of joining social groups is the possibility of social transmission of information related to novel behaviours or resources. Network-based diffusion analysis (NBDA) has emerged as a leading frequentist framework for inferring and quantifying social transmission, particularly in non-human animal populations.
2. NBDA has been extended several times to account for multiple diffusions, multiple networks, individual-level variables, and complex transmission functions. Bayesian versions of NBDA have been proposed before, although these implementations have seen limited usage and have not kept pace with the evolving ecosystem of Bayesian methods. There is not yet a user-friendly package to implement a Bayesian NBDA.
3. Here, we present a unified framework for performing Bayesian analysis of social transmission using NBDA-type models, implemented in the widely used Stan programming language. We provide a user-friendly R package “STbayes” (ST: social transmission) for other researchers to easily use this framework. STbayes accepts user-formatted data, but can also import data directly from the existing NBDA R package. Based on the data users provide, STbayes automatically generates multi-network, multi-diffusion models that allow for covariates that may influence transmission, and varying (random) effects.
4. Using simulated data, we demonstrate that this model can accurately differentiate the relative contribution of individual and social learning in the spread of information through networked populations. We illustrate how incorporating upstream uncertainty about the relationships between individuals can improve model fit. Our framework can be used to infer complex transmission rules, and we describe a numerically stable parameterization of frequency-dependent transmission. Finally, we introduce support for dynamic transmission weights and a “high-resolution” data mode, which allows users to make use of fine-scale data collected by contemporary automated tracking methods. These extensions increase the set of contexts that this type of model may be used for.

## 1 Introduction

Social interaction allows for the transmission of knowledge, behaviours, states, and diseases between individuals. Transmission is a fundamental process that results in a variety of phenomena in collective behaviour and evolution [1, 2, 3]. Identifying and quantifying social transmission, relevant interaction networks, and transmission mechanisms are key questions for researchers interested in social behaviour. Network-based diffusion analysis (NBDA) has emerged as a leading statistical framework for detecting social transmission [4, 5, 1, 6]. The NBDA framework describes a family of time-to-event models. Time-to-event models, such as survival models, typically estimate hazard rates from distributions of deaths over a given observation period. Rather than mortality events, NBDA has been used to study acquisition events where individuals acquire a novel, potentially socially learned behaviour. By combining data about the orders or times of events with the network structure of the population, NBDA can be used to infer the extent to which social transmission plays a role in the spread of a behaviour. NBDA assumes that if social transmission is occurring, it follows the provided network: naive individuals that are well-connected to knowledgeable models are expected to learn sooner than less well-connected individuals. However, these types of model could be applied in other contexts, such as disease or information transmission, and for the remainder of the paper we use the term “event” as a catch-all for any of these contexts.

NBDA models minimally require network data and event data where each observation is the occurrence of an event. In a social learning context, events are often operationalized as the first observation of the usage of the target behaviour by each individual. The level of detail of either data-type determines the specification of the log-likelihood. If only the order of events is known, one can perform an order of acquisition diffusion analysis (OADA), where for each event, the likelihood specifies the probability of the next event occurring to individual *i* amongst the set of all other individuals for which the event has not occurred. If the times of events are known, one can perform a time of acquisition diffusion analysis (TADA). Different to OADA, the likelihood is specified by the joint probability of the event not happening to individual *i* before time *t*, and an event happening to individual *i* at time *t*.

Since it’s introduction in 2009 [4, 5], NBDA has been used to identify social transmission in a variety of taxa. Researchers have used it to determine whether or not novel behaviours were being socially learned in primates [7, 8, 9, 10, 11], cetaceans [12, 13, 14], fish [15, 16], birds [17, 18, 19, 20, 21], and honey-bees [22]. NBDA has been adapted and extended to consider multiple networks [23], dynamic networks where nodes change their connections over time [8], and complex contagion, such as frequency dependent rules for acquisition [24, 25].

Bayesian implementations have been introduced before, in the context of spatial diffusions [26, 27], applying varying effects [28, 22] and Bayesian model comparison [29]. However, most of these studies were published a decade ago, and the ecosystem of popular software and methods for fitting Bayesian models has evolved in the interim period. For example, Stan [30] has emerged as a dominant framework across fields due to its active online forums and inclusion in popular contemporary textbooks [31, 32, 33, 34]. Bayesian NBDA was developed prior to the emergence of Stan’s popularity, and has seen limited use or further development. To date, there is no unified framework for creating, fitting and comparing Bayesian models of social transmission.

Here, we present a unified, user-friendly framework for performing Bayesian analysis of social transmission using NBDA-type models implemented in Stan. We create an R package “STbayes” that serves as a familiar interface between users and Stan, and greatly simplifies the workflow for organizing data, generating models, fitting those models, interpreting outputs and inference via model comparison. Additionally, we present several extensions of the NBDA framework. In the sections below we provide a walk-through of the analysis pipeline using STbayes, and introduce our extensions to the framework. Supplementary text S1 provides primer on Bayesian inference in the context of social transmission, and table S1 gives a summary of notation.

## 2 The pipeline from data to inference in STbayes

The core functionality of STbayes is to 1) facilitate the intake and organization of data, 2) generate specifications of models in Stan based on that data and a users needs, 3) compile and fit those models, and 4) interpret the output of the models. STbayes implements implements several extensions of the framework, including the propagation of uncertainty from network models to transmission models, the use of cross-validation methods for model comparison, the inclusion of random effects on any parameter, time-varying transmission weights, posterior distributions of estimated event times, a novel parameterization of frequency-dependent biases, and usage of very fine-scale data generated by new experimental methodologies. We first provide a step-by-step overview of how to use the full pipeline of STbayes, and then describe each of the novel additions in turn, validated with simulated data. We direct readers to the vignettes available in the Data Availability section for more details and examples.

### 2.1 Data intake

Minimally, STbayes needs an event and a network dataframe, illustrated in Figure 1A. The event dataframe indicates the time at (or order in) which events occurred. The network dataframe is an edge list defining connections between dyads. Networks can be weighted or binary, directed or undirected, symmetric (all combinations of individuals provided, useful for directed networks) or asymmetric. Supplying multiple networks will generate a multi-network model that estimates the strength of social transmission for each network. For an in-depth discussion of the interpretation of results under different network types, we direct readers to Hoppitt 2017 [35]. Users may give dynamic networks that change over the observation period. If users already use the NBDA package, STbayes also accepts NBDA objects.

**Figure 1.**
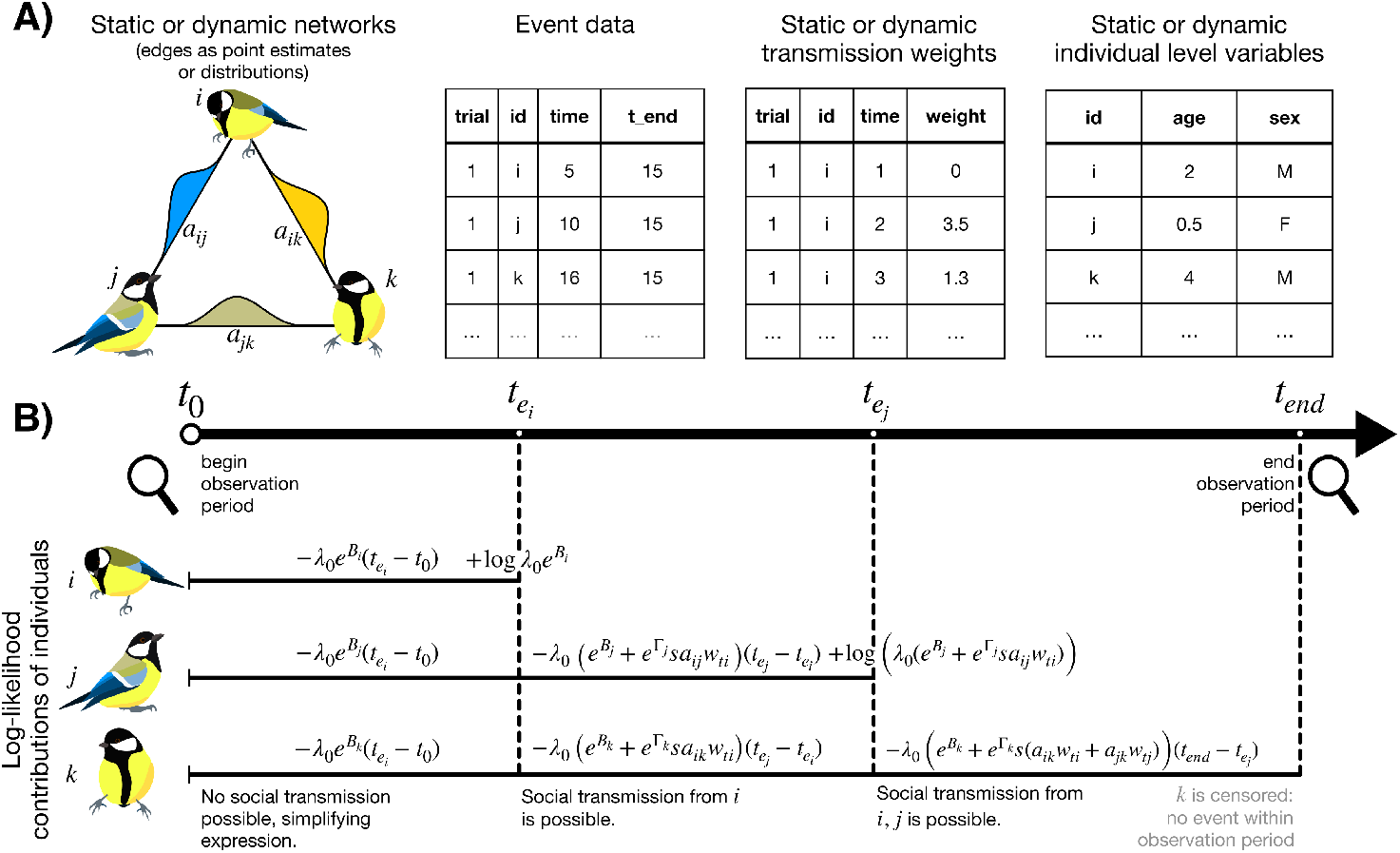
STbayes overview. A) Summary of data types. STbayes requires network data that defines edge-weights between individuals, and event data, either the order in which or times that events occurred. Optionally, users can supply static or dynamic transmission weights, representing the overall rate, or time-varying rates of behavioural usage or relevant cue production. Users may also provide static or time-varying ILVs. B) Toy example of log-likelihood function for a TADA-type model. In this example, three individuals are under observation, and individuals *i* and *j* acquire the target behaviour within the observation period. Shown are individuals’ contributions to the log-likelihood during each inter-event period, as well as the contribution of when an event occurs for *i* and *j*.

Continuous or categorical individual-level variables (ILVs) may also be input. Constant ILVs can be included using a dataframe with identities and values of one or more variable. Time-varying ILVs require an additional time column, and like the networks dataframe above, variables should be summarized for inter-event periods. ILVs can be treated additively (separate effects on the intrinsic and social rates) or multiplicatively (single effect on both rates).

### 2.2 Model generation

Next, STbayes dynamically generates an appropriately specified Stan model using a single function call. Users can specify whether the times of events are known (creating a TADA type model) or only the order of events is known (OADA type model). If exact times are known, users should use a cTADA model. A dTADA model is appropriate if a population was regularly (or irregularly) sampled, so the time of events are unknown but the rough interval they occurred in is known. If order, rather than time, of events is known, users can specify an OADA model, whose likelihood is described in supplementary text S2. Further discussion of the distinctions between these models, as well as a decision tree for choosing the appropriate model is provided in Hasenjager, Leadbeater, and Hoppitt [6].

Users may also specify 1) whether to generate a full model (both social and intrinsic rates) or an asocial model (intrinsic rate only), 2) use a constant intrinsic rate, or a Weibull shaped rate that can increase or decrease over time, 3) use simple or complex transmission, 4) whether to include a generated quantities block to output the log-likelihood of observations for model comparison, 5) whether to output estimated acquisition times in the generated quantities block, useful for posterior predictive checks, and 6) the priors used (see Figure S1 and supplementary text S3 for a prior predictive check and further details). Finally, if the user is modelling repeated trials with the same individuals, it is possible to fit varying effects (random effects in frequentist terminology) on individuals, trials or both individuals and trials. Advanced users can easily save the Stan model for further modification.

### 2.3 Model fitting

Users can then pass the data and the model to a single fitting function that interfaces with CmdStanR [36]. The model will compile and run, and the function returns an object of class CmdStanMCMC. Users can pass any other valid arguments to CmdStanR, such as the number of chains and iterations, and other control parameters. Default settings (1 chain, 1000 iterations) are enough to give an impression of the posterior while minimizing the fitting time, but users should run ≥ 4 chains to accurately sample the posteriors’ tails. We give further guidance in supplementary text S4. Simple models will take seconds to fit, but time increases depending on hardware, model complexity, amount of data, number of iterations, and hyper-parameters. Runtimes under different amounts of data are summarized in Figure S4.

We provide a function that saves the fit and associated csv files containing samples to a named RDS file. By default, CmdStanR will save these csvs to a temporary folder, which is purged when R is restarted. If users are not familiar with CmdStanR, we recommend using this function.

### 2.4 Interpreting data and model comparison using cross-validation methods

If the user is familiar with cmdstanR, they may proceed to use their own analysis pipeline. For beginners, STbayes provides convenient functionality for understanding model outputs. Users can output a summary table of parameter estimates, confidence intervals, and diagnostics for parameters. We provide an example output from a single-network model in Table S2.

Of most importance are estimates of the intrinsic rate (*λ*_0_, previously referred to as the asocial learning rate) and the relative strength of social transmission (*s*). *λ*_0_ represents the underlying rate of a target event in the absence of social transmission. *s* represents the rate of transmission per unit connection relative to the intrinsic rate. For a detailed description of these parameters and their context in the likelihood, please see supplementary text S1.

STbayes also calculates the percentage of events that occur through social transmission (%ST). The probability of event *e* occurring through social transmission is the social component of the hazard rate divided by the full rate (intrinsic and social component) at the time of event. %ST is then the mean probability over all events. This can be generalized to multi-network models by calculating the probability that each event occurred through social transmission on network *n* (see supplementary text S5). The NBDA package produced confidence intervals of %ST by re-calculating this value using the upper and lower 95% CI for *s* [6]. STbayes refines this by computing the posterior distribution of %ST in the generated quantities block of the STbayes model, which includes full uncertainty over all parameters.

For model comparison, we provide a convenient function to compare multiple models using functions from the *loo* package [37]. The NBDA package uses AIC (Akaike information criterion) for model comparison [38], using Δ*AIC* between constrained asocial models and social transmission models as evidence for social transmission [6]. However, there are several more contemporary options available for Bayesian model comparison using existing cross-validation tools, such as WAIC (Widely Applicable Information Criterion) [39] or LOO-PSIS (leave-one-out via pareto smoothed importance sampling) cross-validation [37], both of which use in-sample fits to estimate out-of-sample predictive accuracy. AIC assumes a single set of point estimates for parameter values that maximizes the likelihood function (MLE), and ignores any uncertainty around estimates. In comparison, WAIC and LOO-PSIS integrate over the entire posterior, accounting for uncertainty. While it is beyond the scope of this paper to fully describe cross validation, we provide a basic description in supplementary text S6.

## 3 Validation and illustrative examples

In order to validate that STbayes performs similarly to the frequentist NBDA package, we conducted 100 stimulated diffusions with known parameter values and fit models to the same data using both packages. See supplementary text S7 for a description of simulations. Estimates were comparable (Figure S2) and also strongly correlated between packages (Figure S3).

In order to provide guidance on study design, we performed a power analysis using networks of different population sizes, densities, and number of trials, detailed in supplementary text S8 and figure S5. A vignette on performing power analyses is provided in the online documentation. We found that given a modest effect of social transmission (*s* = 2), so long as the study population is large enough (≥ 24), a single trial is enough to reliably support social transmission. We validated that STbayes can recover varying effects of individuals (Figure S6), although the accuracy of the estimates decreases with fewer trials (Figure S7). If researchers wish to estimate varying effects of individuals, we suggest using ≥ 50 trials.

### 3.1 Accounting for network uncertainty

Social networks are inferred from noisy observations, meaning that there is inherent uncertainty in edge-weights. One recommendation has been to use a threshold rule, removing individuals with few observations from social networks [40, 41]. However, missing edges can lead to the inference that an event occurred due to the intrinsic rate, rather than social transmission. Wild and Hoppitt [42] provided a tool for determining an appropriate threshold that would not be overly-conservative for NBDA models. Recent Bayesian tools, such as the bisonR package [43] or STRAND package [44, 45] estimate edge-weights as distributions that reflect uncertainty, circumventing the need for using a threshold. STbayes can account for uncertainty in network measurement by feeding forward the posterior distribution of edge-weights from model fits using bisonr or STRAND (or lists of model fits in the case of multi-network models). Following the approach of Hart et al. [43], the model includes edge-weights modelled by a multivariate normal approximation using the sample mean and covariance matrix calculated from the posterior edge-weight samples. This allows the joint uncertainty over edge-weights to be accounted for.

To evaluate the impact of accounting for network uncertainty in social transmission models, we 1) simulated binary interaction data from a latent network with uneven sampling effort for different dyads, 2) fit a Bayesian network model using bisonr [43] to infer posterior edge distributions, and then 3) simulated 50 diffusion trials using the true network with known parameter values. We fit two transmission models to the resulting event data: one where edge-weights were point estimates (the median value of edges from the network model), and another where edge-weights were posterior distributions from the network model. Both models reasonably recovered the overall hazard (Figure 2A). The point estimate model over-estimated *λ*_0_ (Table S3), while the model that marginalized over uncertainty recovered *λ*_0_ accurately (Table S4). It better recovered the relative relationship between intrinsic and social rates (Figure 2B) and obtained significantly better predictive performance (LOOIC = 24520.6) than the point-estimate model (LOOIC = 24636.3). In summary, marginalizing over the posteriors of edges can help prevent the model from confidently over-fitting the data, especially in cases where there is uneven sampling effort among individuals.

**Figure 2.**
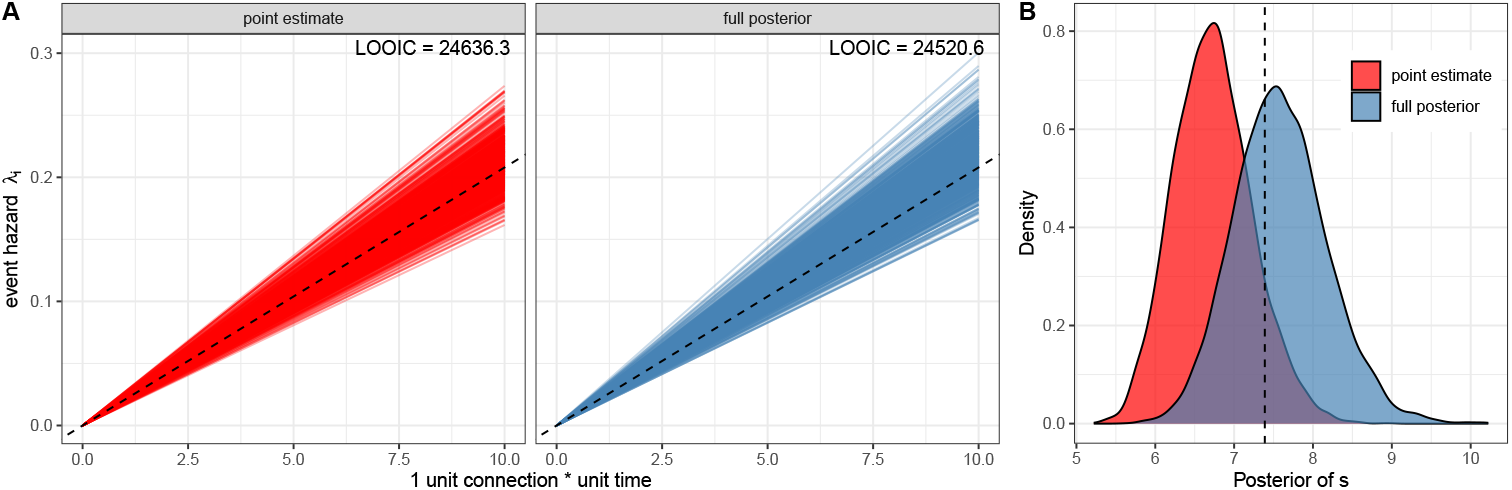
Estimated relationship between number of informed associates and hazard rate. STbayes allows for the propagation of uncertainty from network models to the estimation of diffusion parameters. A) We compared a model fitted with networks as point estimates (calculated as a simple ratio index, left panel) against a model incorporating uncertainty from the network model (right panel). Dashed lines indicate the ground truth hazard function used to simulate the data. Incorporating uncertainty in network edges can provide a more accurate estimate of parameters with better out-of-sample predictions as measured by LOOIC. B) Marginalizing over uncertainty can improve the estimate of the relative strength of social transmission.

There may be other types of measurement uncertainty in ILVs or missing data. Due to the complexity of automated model generation, it is difficult to create a generalisable solution in STbayes itself. However, advanced users can save and directly modify the Stan code for such cases, and pass the file path of the modified model to the fitting function.

### 3.2 Complex transmission rules

STbayes also allows for customized likelihoods that represent non-linear “complex” transmission mechanisms, such as frequency-dependent transmission [24, 25]. The user can choose among standard linear transmission, and two options of complex transmission, which we refer to as “complex f” and “complex k”. “Complex f” is the form traditionally used in social learning studies [46, 24]. We introduce “complex k” as an alternative parameterization. “Complex k” improves on “complex f” because 1) it is more numerically stable during sampling: fitting *k* requires sampling a narrow range of values, and there is no exponentiation, and 2) the effect of *k* on the shape of the curve is symmetrical whereas *f* is asymmetrical. Supplementary text S9 details each. Both have a similar effect on how proportions of associates are converted into transmission weights (Figure 3). We note that complex transmission changes the interpretation of *s*, as it is no longer relative to a unit connection to a knowledgable individual. Rather, *s* is relative to when *all* connections are with knowledgable individuals. If *s* = 2, under standard transmission individual *i* is twice as likely to learn a behaviour if *a*_*ij*_ = 1 and *j* is knowledgable. Under complex transmission, they are twice as likely to learn a behaviour if 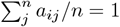 and all *n* neighbours are knowledgeable.

**Figure 3.**
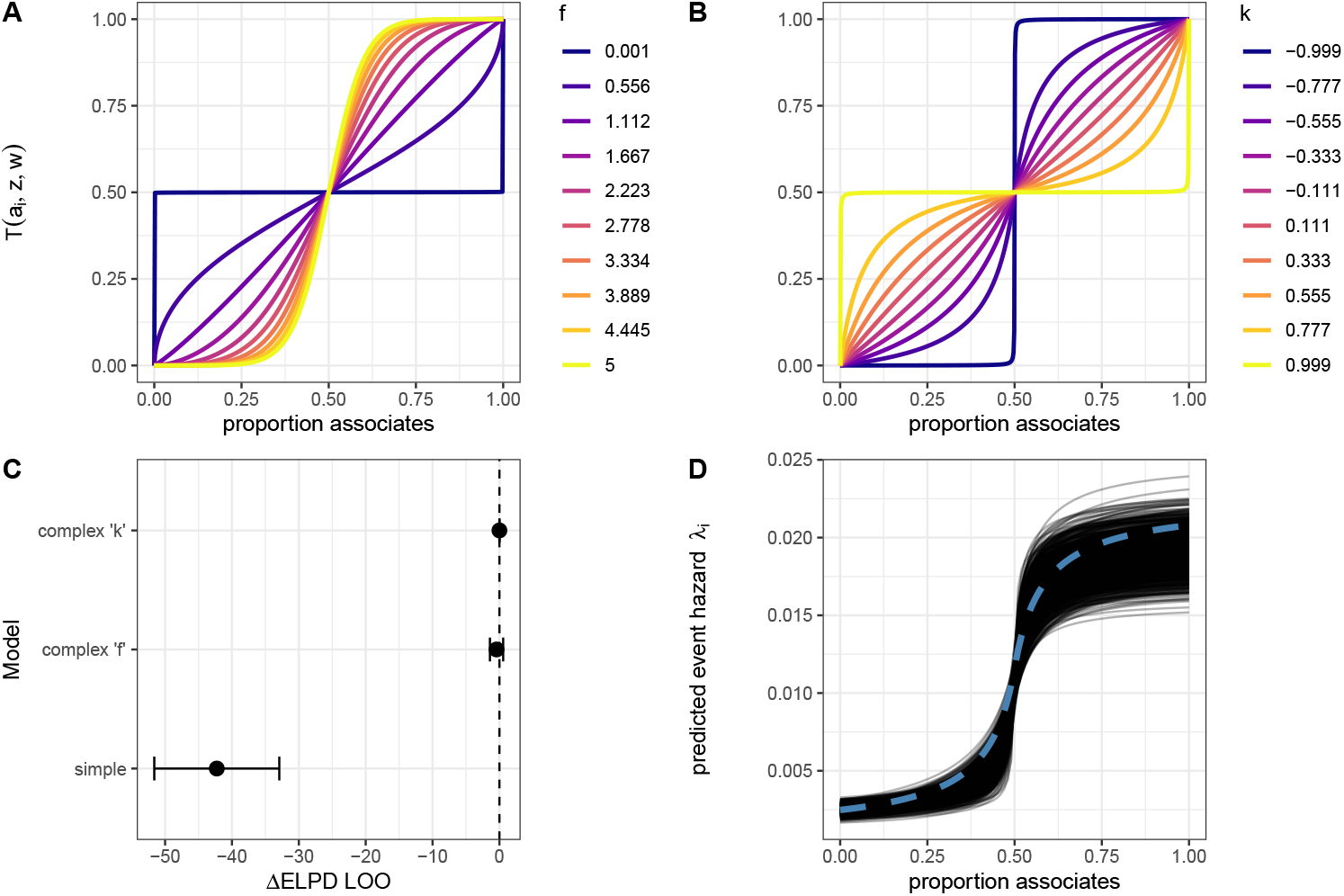
Alternative parameterizations of complex transmission. A) Traditional frequency-depended formulation with *f* parameter. B) Normalized tuneable sigmoid with parameter *k*. Both parameterizations result in similar transformations and interpretations of the shape parameters. C) Both parameterizations result in similarly good predictive performance as measured by differences in Δ*ELPD*_*loo*_ on simulated data. D) Estimated hazard as the proportion of informed associates increases from the complex k parameterization.

We validated both complex models with simulated data from 10 trials of 50 individuals each with known parameter values using static networks. Both parameterizations supported significant terms for positive frequency dependent biases, and Δ*ELPD*_*loo*_ showed that they were better fit to the data than a model of simple transmission, but there was no significant difference in predictive performance between the two parameterizations (Figure 3C, D). We strongly suggest that researchers use multiple trials (≥ 10 trials), as fits of individual trials produced wide error margins for *k* and *f* estimates. Sampling efficiency was poorer with “complex f”: a model with 5 chains of 5000 iterations had a mean chain execution time of 519.1 seconds, and 200 seconds under “complex k”.

### 3.3 Dynamic networks and transmission weights

The NBDA package accepts static transmission weights (*w*_*i*_) that represent a summary rate of production or usage of the target behaviour by an individual over the observation period. STbayes extends this to allow for dynamic transmission weights (*w*_*it*_) if researchers have detailed behavioural time-series data. This is important for accounting for variable behavioural production rates (in the context of social learning of a novel behaviour) or variable shedding rates (in the context of pathogen transmission) [47]. For example, if an individual knows a target behaviour, but doesn’t use a target behaviour during an inter-event interval, then its influence on others should be null during that time period.

To illustrate the value of including both dynamic networks and dynamic transmission weights, we simulated diffusion data from a spatially explicit, motile population of learners similar to the approach of Chimento and Farine [25]. The details of these simulations may be found in supplementary text S7. We recorded their interaction network each timestep (dynamic network *A*_*ijt*_, Figure 4A), and an overall summary of their interaction for the duration of the observation period (static network *A*_*ij*_, Figure 4B). We recorded their usage of the target behaviour over time (dynamic transmission weight *w*_*it*_), and a summary for the duration of the observation period (static transmission weight *w*_*i*_). We then fit incrementally more complex models, illustrating that inclusion of all relevant data results in a better model fit (Figure 4C,D,E).

**Figure 4.**
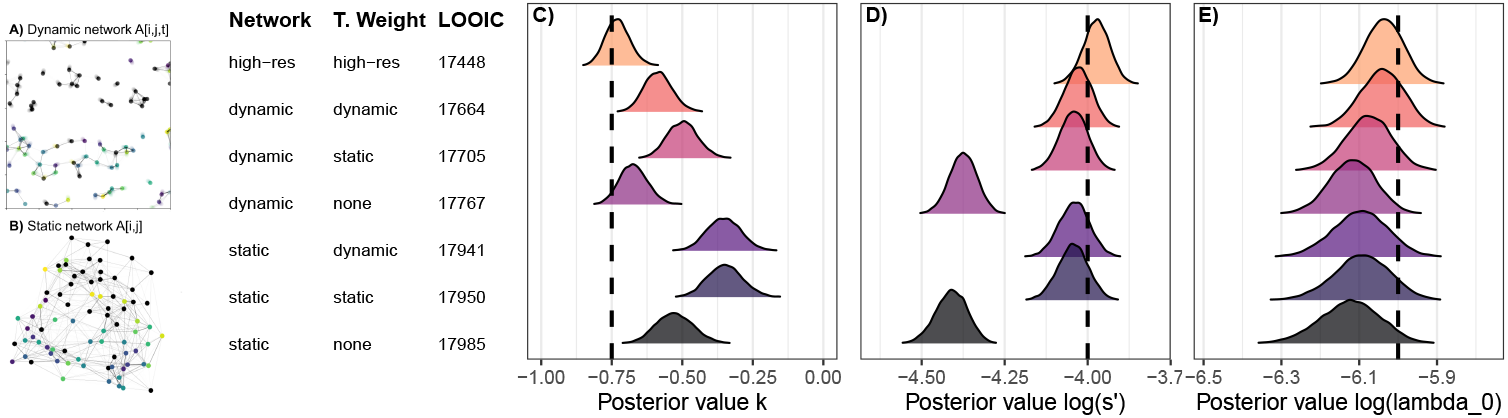
Using dynamic transmission weights and networks can improve model fit and estimates. A) We simulated diffusions using a positive frequency-dependent complex transmission rule on dynamic networks of motile agents, and then B) summarized their network over the duration of the observation period, as would be done to an empirical study. Using these dynamic and static networks, we fit increasingly complex models that contained more information about the diffusion. Posterior distributions of the frequency dependent parameter, log(*s*′) and log(*λ*_0_) are shown in panels C, D and E. Providing dynamic networks and transmission weights improved predictive performance and estimates. “High-resolution” models, which contained dynamic networks and dynamic transmission rates measured at each timestep resulted in the best model fit and most accurate parameter estimates.

### 3.4 “High-resolution” data mode

NBDA models have been limited to incorporating dynamic networks and time-varying ILVs that only change at event cut-points. Above, we introduced the ability to allow transmission weights to change at cut-points. When data was generated using motile agents who could continuously move, model estimates were less accurate than expected due to the omission of fine-scaled detail about the diffusion, as networks and transmission weights were summarized over inter-event intervals. We introduce “high-resolution” data mode, where these values may vary independently of event times. This is useful when network connections and transmission weights are misaligned due to the movement of individuals. While it has been practically difficult to collect high-resolution data throughout the course of a diffusion, recent methodological advances, such as multi-individual tracking have created the opportunity to do so [48, 49, 50].

Complex transmission creates especially difficult conditions to estimate parameters from cumulative representations of networks [25]. Parameter estimates were most accurate when using high resolution mode (Figure 4). We find that providing dynamic transmission weights provides a good improvement in parameter estimates for *s*′, although not for the frequency dependent parameter (Figure 4C,D). However, providing dynamic networks, even without dynamic transmission weights, helps identify the frequency dependent parameter (Figure 4C). We recommend care when interpreting the precise values of *k* or *f* when applied to real datasets in the absence of highly detailed dynamic networks and transmission weights. It is also logical that inference about complex transmission becomes more difficult when the average degree of a network is low, as it might not provide enough information about the shape of the relationship between the unit connection to informed individuals and hazard rates.

We additionally tested the same set of models on data simulated from motile agents using the standard transmission rule. Here, providing dynamic networks or dynamic transmission weights significantly improved model fit and parameter estimates (Figure S8). “High-resolution” mode improved the model fit, but didn’t significantly improve parameter estimates from dynamic networks or dynamic weights models, as they were already quite close to the true values. This might depend on many factors, including the types of movements, length of diffusion, and strength of social transmission. Beyond the case of complex transmission in motile populations, more research is needed to understand when it is critical to include extremely high resolution data.

Fitting a model with complex transmission in high-resolution mode may take a long time. Compared to the tens to several hundreds of seconds for all other models presented in this study, this model specification took 170 minutes (*N* = 10 trials, 50 individuals, 5 parallel chains for 5000 iterations). For standard transmission, STbayes optimizes fitting a dataset of the same size in under 8 minutes.

## 4 Discussion

STbayes introduces a flexible yet user-friendly framework for modelling social transmission using Bayesian inference. While the frequentist NBDA approach has been a valuable tool, it has lacked a unified, user-friendly Bayesian alternative. STbayes fills this gap, allowing users to specify complex models of diffusion processes with fine control over priors, model structure, and uncertainty propagation. By building on the Stan ecosystem, STbayes benefits from contemporary advances in Bayesian computation.

While NBDA models has been largely been applied in the field of social learning and cultural transmission, we emphasize that this family of models can be used in a variety of contexts, as long as there are measurable events and measurable interactions. Our introduction of dynamic transmission weights expands the applicability of these models to recurrent events that are relevant for the study of social contagion: yawning, sleep or other state statuses that may vacillate over time. Dynamic transmission weights also offer researchers creativity in hypothesis testing about which specific socially observed cues influence the hazard rate of a target event. They may perform model comparison between models fit on dynamic transmission weight data of different cues.

Another contribution of STbayes is the ability to propagate uncertainty from network models to transmission models: a meaningful step toward more robust inference in studies of social transmission. Social network data are almost always estimates based on limited observations with unequal sampling effort. STbayes allows users to incorporate posterior distributions from upstream network models (e.g., BISON or STRAND) directly into the inference process, thereby accounting for uncertainty in network structure [43, 44, 45].

Another key extension is the inclusion of non-linear, frequency-dependent transmission functions. These allow researchers to test for frequency-dependent biases in the transmission process. STbayes includes two parameterizations of complex contagion, both of which yield similar predictive performance, though the alternative sigmoid formulation is more numerically stable.

Finally, STbayes encourages a shift away from AIC toward more contemporary Bayesian methods for model comparison, such as LOO-PSIS or WAIC [37, 39]. These methods integrate over the full posterior distribution rather than relying on point estimates, making them more appropriate for potentially sparse and noisy data that might be collected in studies of social transmission.

As researchers move toward more complex and nuanced models of transmission, STbayes offers a platform for inference that is adaptable to a wide range of research questions. We hope that this framework not only improves the reproducibility and robustness of inference, but also encourages further exploration of social transmission processes in both natural and experimental settings.

## 5 Data Availability

Package documentation, including installation instructions and vignettes can be found at https://michaelchimento.github.io/STbayes/index.html. The package will receive an official release until after acceptance. Code and data for statistical analyses and main text figures is available on Edmond [51].

## 6 Author Contributions

Conceptualisation, M.C., W.H.;

Methodology, M.C., W.H.;

Software, M.C.;

Investigation, M.C.;

Visualization, M.C.;

Writing - Original Draft, M.C.;

Writing - Review & Editing, M.C., W.H.

## 7 Acknowledgements

MC received support from the Centre for the Advanced Study of Collective Behaviour, funded by the Deutsche Forschungsgemeinschaft (DFG) under Germany’s Excellence Strategy (EXC 2117-422037984) and the Swiss State Secretariat for Education, Research and Innovation (SERI) under contract number MB22.00056. The Max Planck Society provided open access data storage.

## References

[1] William Hoppitt and Kevin N Laland. Social learning: an introduction to mechanisms, methods, and models. Princeton University Press, 2013.

[2] Mauricio Cantor et al. “The importance of individual-to-society feedbacks in animal ecology and evolution”. In: Journal of Animal Ecology 90.1 (2021), pp. 27–44.

[3] Christos C Ioannou and Kate L Laskowski. “A multi-scale review of the dynamics of collective behaviour: from rapid responses to ontogeny and evolution”. In: Philosophical transactions of the royal society B 378.1874 (2023), p. 20220059.

[4] Mathias Franz and Charles L Nunn. “Network-based diffusion analysis: a new method for detecting social learning”. In: Proceedings of the Royal Society B: Biological Sciences 276.1663 (2009), pp. 1829–1836.

[5] William Hoppitt, Neeltje J Boogert, and Kevin N Laland. “Detecting social transmission in networks”. In: Journal of Theoretical Biology 263.4 (2010), pp. 544–555.

[6] Matthew J Hasenjager, Ellouise Leadbeater, and William Hoppitt. “Detecting and quantifying social transmission using network-based diffusion analysis”. In: Journal of Animal Ecology 90.1 (2021), pp. 8–26.

[7] Rachel L Kendal et al. “Evidence for social learning in wild lemurs (Lemur catta)”. In: Learning & Behavior 38.3 (2010), pp. 220–234.

[8] Catherine Hobaiter et al. “Social network analysis shows direct evidence for social transmission of tool use in wild chimpanzees”. In: PLoS Biol 12.9 (2014), e1001960.

[9] Stuart K Watson et al. “Socially transmitted diffusion of a novel behavior from subordinate chimpanzees”. In: American journal of primatology 79.6 (2017), e22642.

[10] Charlotte Canteloup, William Hoppitt, and Erica van de Waal. “Wild primates copy higher-ranked individuals in a social transmission experiment”. In: Nature communications 11.1 (2020), pp. 1–10.

[11] Camila Galheigo Coelho et al. “Social tolerance and success-biased social learning underlie the cultural transmission of an induced extractive foraging tradition in a wild tool-using primate”. In: Proceedings of the National Academy of Sciences 121.48 (2024), e2322884121.

[12] Jenny Allen et al. “Network-based diffusion analysis reveals cultural transmission of lobtail feeding in humpback whales”. In: Science 340.6131 (2013), pp. 485–488.

[13] Sonja Wild et al. “Multi-network-based diffusion analysis reveals vertical cultural transmission of sponge tool use within dolphin matrilines”. In: Biology letters 15.7 (2019), p. 20190227.

[14] Sonja Wild et al. “Integrating genetic, environmental, and social networks to reveal transmission pathways of a dolphin foraging innovation”. In: Current Biology 30.15 (2020), pp. 3024–3030.

[15] Mike M Webster et al. “Environmental complexity influences association network structure and network-based diffusion of foraging information in fish shoals”. In: The American Naturalist 181.2 (2013), pp. 235–244.

[16] Matthew J Hasenjager, William Hoppitt, and Lee A Dugatkin. “Personality composition determines social learning pathways within shoaling fish”. In: Proceedings of the Royal Society B 287.1936 (2020), p. 20201871.

[17] Damien R Farine, Karen A Spencer, and Neeltje J Boogert. “Early-life stress triggers juvenile zebra finches to switch social learning strategies”. In: Current Biology 25.16 (2015), pp. 2184–2188.

[18] Virginia K Heinen et al. “Food discovery is associated with different reliance on social learning and lower cognitive flexibility across environments in a food-caching bird”. In: Proceedings of the Royal Society B 288.1951 (2021), p. 20202843.

[19] Barbara C Klump et al. “Innovation and geographic spread of a complex foraging culture in an urban parrot”. In: Science 373.6553 (2021), pp. 456–460.

[20] Ipek G Kulahci et al. “Social networks predict selective observation and information spread in ravens”. In: Royal Society open science 3.7 (2016), p. 160256.

[21] Zoltán Tóth et al. “The effect of social connections on the discovery of multiple hidden food patches in a bird species”. In: Scientific reports 7.1 (2017), pp. 1–9.

[22] Matthew J Hasenjager et al. “Coupled information networks drive honeybee (Apis mellifera) collective foraging”. In: Journal of Animal Ecology 93.1 (2024), pp. 71–82.

[23] Damien R Farine et al. “Interspecific social networks promote information transmission in wild songbirds”. In: Proceedings of the Royal Society B: Biological Sciences 282.1803 (2015), p. 20142804.

[24] Josh A Firth et al. “Analysing the Social Spread of Behaviour: Integrating Complex Contagions into Network Based Diffusions”. In: arXiv preprint arXiv:2012.08925 (2020).

[25] Michael Chimento and Damien R Farine. “The contribution of movement to social network structure and spreading dynamics under simple and complex transmission”. In: Philosophical Transactions B 379.1912 (2024), p. 20220524.

[26] Glenna F Nightingale et al. “Bayesian spatial NBDA for diffusion data with home-base coordinates”. In: PLoS One 10.7 (2015), e0130326.

[27] Glenna Nightingale et al. “Quantifying diffusion in social networks: a Bayesian approach”. In: Animal social networks (2015), pp. 38–52.

[28] Neeltje J Boogert et al. “Perching but not foraging networks predict the spread of novel foraging skills in starlings”. In: Behavioural processes 109 (2014), pp. 135–144.

[29] Andrew Whalen and William JE Hoppitt. “Bayesian model selection with network based diffusion analysis”. In: Frontiers in psychology 7 (2016), p. 409.

[30] Bob Carpenter et al. “Stan: A probabilistic programming language”. In: Journal of statistical software 76 (2017), pp. 1–32.

[31] Andrew Gelman et al. Bayesian Data Analysis. CRC Press, 2013.

[32] Franzi Korner-Nievergelt et al. Bayesian data analysis in ecology using linear models with R, BUGS, and Stan. Academic Press, 2015.

[33] Richard McElreath. Statistical rethinking: A Bayesian course with examples in R and Stan. Chapman and Hall/CRC, 2018.

[34] Alicia A Johnson, Miles Q Ott, and Mine Dogucu. Bayes rules!: An introduction to applied Bayesian modeling. Chapman and Hall/CRC, 2022.

[35] Will Hoppitt. “The conceptual foundations of network-based diffusion analysis: choosing networks and interpreting results”. In: Philosophical Transactions of the Royal Society B: Biological Sciences 372.1735 (2017), p. 20160418.

[36] Jonah Gabry et al. cmdstanr: R Interface to ‘CmdStan’. R package version 0.9.0, https://discourse.mc-stan.org. 2025. xurl: https://mc-stan.org/cmdstanr/.

[37] Aki Vehtari, Andrew Gelman, and Jonah Gabry. “Practical Bayesian model evaluation using leave-one-out cross-validation and WAIC”. In: Statistics and computing 27 (2017), pp. 1413–1432.

[38] Hirotogu Akaike. “Information theory and an extension of the maximum likelihood principle”. In: ed. By F. Caski B.N. Petrov. Akademiai Kiado, 1973, pp. 267–281.

[39] Sumio Watanabe and Manfred Opper. “Asymptotic equivalence of Bayes cross validation and widely applicable information criterion in singular learning theory.” In: Journal of machine learning research 11.12 (2010).

[40] Daniel W Franks, Graeme D Ruxton, and Richard James. “Sampling animal association networks with the gambit of the group”. In: Behavioral ecology and sociobiology 64 (2010), pp. 493–503.

[41] Matthew J Silk et al. “The consequences of unidentifiable individuals for the analysis of an animal social network”. In: Animal Behaviour 104 (2015), pp. 1–11.

[42] Sonja Wild and William Hoppitt. “Choosing a sensible cut-off point: assessing the impact of uncertainty in a social network on the performance of NBDA”. In: Primates 60.3 (2019), pp. 307–315.

[43] Jordan Hart et al. “BISoN: a Bayesian framework for inference of social networks”. In: Methods in ecology and evolution 14.9 (2023), pp. 2411–2420.

[44] Daniel Redhead, Richard McElreath, and Cody T Ross. “Reliable network inference from unreliable data: A tutorial on latent network modeling using STRAND.” In: Psychological methods (2023).

[45] Cody T Ross, Richard McElreath, and Daniel Redhead. “Modelling animal network data in R using STRAND”. In: Journal of Animal Ecology 93.3 (2024), pp. 254–266.

[46] Richard McElreath et al. “Beyond existence and aiming outside the laboratory: estimating frequency-dependent and pay-off-biased social learning strategies”. In: Philosophical Transactions of the Royal Society B: Biological Sciences 363.1509 (2008), pp. 3515–3528.

[47] Michael Chimento et al. “Cultural diffusion dynamics depend on behavioural production rules”. In: Proceedings of the Royal Society B 289.1980 (2022), p. 20221001.

[48] Tristan Walter and Iain D Couzin. “TRex, a fast multi-animal tracking system with markerless identification, and 2D estimation of posture and visual fields”. In: eLife 10 (Feb. 2021). Ed. by David Lentink, e64000. issn: 2050-084X. doi: 10.7554/eLife.64000. url: https://doi.org/10.7554/eLife.64000.

[49] Máté Nagy et al. “SMART-BARN: Scalable multimodal arena for real-time tracking behavior of animals in large numbers”. In: Science Advances 9.35 (2023), eadf8068.

[50] Mathilde Delacoux and Fumihiro Kano. “Fine-scale tracking reveals visual field use for predator detection and escape in collective foraging of pigeon flocks”. In: Elife 13 (2024), RP95549.

[51] Michael Chimento and William Hoppitt. Data and Code to Reproduce “STbayes: An R package for creating, fitting and understanding Bayesian models of social transmission”. Version V2. 2025. doi: 10.17617/3.IY3TKX. url: https://doi.org/10.17617/3.IY3TKX.

