## supplementary for "STbayes: An R package for creating, fitting and understanding Bayesian models of social transmission"

#### Supplementary Materials

#### Text

- **Text S1**: Bayesian inference in a social transmission context
- **Text S2**: Order of acquisition diffusion analysis (OADA) likelihood
- **Text S3**: Definition of priors
- **Text S4**: Guidance on MCMC model fitting
- **Text S5**: Definition of %ST
- **Text S6**: Description of cross-validation model comparison
- **Text S7**: Description of simulations used for model validation
- **Text S8**: Description of power analysis
- **Text S9**: Expanded description of WAIC and LOO-PSIS.

#### Figures

- **Figure S1**: Prior predictive check.
- **Figure S2**: Comparison of estimated parameters between NBDA and STbayes
- **Figure S3**: Correlation between NBDA and STbayes estimates.
- **Figure S4**: Model runtimes
- **Figure S5**: Power analysis results
- **Figure S6**: STbayes estimates of main and varying effects.
- **Figure S7**: Varying effects estimates with lower sample sizes.
- **Figure S8**: Testing dynamic transmission weights and networks under simple transmission.

#### Tables

- **Table S1**: Summary of notation
- **Table S2**: Typical output from STb_summary() function
- **Table S3**: Summary of posterior estimates for model where edges are point estimates (Section 5).
- **Table S4**: Summary of posterior estimates for model where edge uncertainty is included (Section 5).
- **Table S5**: Summary of posterior estimates for complex transmission using a static network (Section 6).

### S1 Bayesian inference in a social transmission context

Bayesian inference can be summarized using Bayes' Rule:

$$P(\theta|D) = \frac{P(D|\theta)P(\theta)}{P(D)}, \tag{1}$$

where $\theta$ are parameters from the data generating process and $D$ is the observed data. $P(D|\theta)$ is the likelihood, or how likely the observed data is given some parameter values, $P(D)$ is the evidence (or marginal likelihood), which is the total probability of data across all possible parameter values, and $P(\theta|D)$ is the posterior, or an updated estimate of parameter values after seeing the data.

When studying social transmission, we are primarily interested in estimating two parameters: an intrinsic event rate ($\lambda_0$) that operates independently of social information, and the strength of social transmission ($s$), which quantifies the influence that interactions with others has on event rates. NBDA has been extended to include additive individual-level variables (ILVs) that might affect either the intrinsic rate or strength of social transmission, or multiplicative ILVs that equally affect both rates [1]. The coefficients of which may written as $\beta$, $\gamma$, and $\phi$ respectively. Thus, in a social transmission context:

$$\theta = \{\lambda_0, s, \beta, \gamma, \phi\}. \tag{2}$$

The data we usually collect in transmission studies are the states of individuals in a population over time ($z \in \{0,1\}$), e.g. whether an event happened yet, such as the acquisition of novel behaviour or pathogen), network connections between individuals $a$, other relevant individual level variables $x$, and ideally transmission weights ($w$, e.g. behavioural production rates, in the context of the spread of a novel behaviour). Thus,

$$D = \{z, a, x, w\}. \tag{3}$$

The likelihood function for given parameter values is composed of two parts: the probability that each individual remained $z_i = 0$ until time $t_e$, and the probability that an individual became $z_i = 1$ at time $t_e$. The likelihood contribution of an individual who experienced an event during the observation period ($i \in N_e$) at time $t_e$ is:

$$\log \mathcal{L}_{\text{uncensored}i} = -\sum_{t_0}^{t_e} \lambda_{it} + \log \lambda_{it_e}. \tag{4}$$

For individuals who do not learn ($i \in N_c$), the likelihood contribution is:

$$\log \mathcal{L}_{\text{censored},i} = -\sum_{t_0}^{t_{\text{end}}} \lambda_{ti}. \tag{5}$$

Here, $\lambda_{ti}$ is the hazard function for individual at time $t$ is given by:

$$\lambda_{ti} = \lambda_0(1 - z_{ti})e^{\Psi_i}\left(e^{B_i} + e^{\Gamma_i} sT(a,z,w,t)\right). \tag{6}$$

In a standard model, the transmission function $T$ defines an simple linear relationship between connections to informed individuals and the hazard rate:

$$T(a,z,w,t) = \sum_j a_{ijt} z_{jt} w_{jt}, \tag{7}$$

although other complex transmission functions may be defined [2, 3], and we have included support for frequency-dependent transmission, detailed in Section 4.2 of the main text. Additive ILVs for the intrinsic rate and social transmission are defined as $B_i = \sum_v \beta_v x_{vi}$ and $\Gamma_i = \sum_v \gamma_v x_{vi}$, respectively. These variable act independently on either rate. Multiplicative ILVs are defined as $\Psi_i = \sum_v \psi_v x_{vi}$. We note that we do not fit $s$ directly, but rather we reparameterize equation 6 to:

$$\lambda_{ti} = (1 - z_{ti})e^{\Psi_i}\left(\lambda_0 e^{B_i} + e^{\Gamma_i} s' T(a,z,w)\right), \tag{8}$$

$s'$ represents $\lambda_0 s$, and the original interpretation of $s$ is recovered by $s'/\lambda_0$ [4]. Both parameters are fit on the log scale for sampling performance, and the original exponentiated interpretations are output by STbayes for easy interpretation. The overall likelihood function can be summarized as:

$$\log \mathcal{L}(\theta|D) = \sum_{k=1}^{K}\left(\sum_{i}^{N_e}\left(-\sum_{t_0}^{t_e}\lambda_{kit} + \log\lambda_{kit_e}\right) + \sum_{i}^{N_{c_k}}\left(-\sum_{t_0}^{t_e}\lambda_{kti}\right)\right). \tag{9}$$

Summation over $K$ diffusions, or trials, allows for multiple diffusions to be included.

STbayes uses the Stan programming language to write and fit models. To sample from the posterior, Stan uses Hamiltonian Monte Carlo (HMC), an advanced Markov Chain Monte Carlo (MCMC) method. HMC uses gradient-based exploration to efficiently sample the probability space. It treats the posterior landscape like a physical system, where a particle moves according to the shape of the probability distribution, allowing it to reach high-probability regions faster and reduce sampling inefficiencies. Importantly, Stan readily accepts customized likelihood functions, making it possible to create flexible, complex models often required for studies of social transmission. Once a model is specified, it is compiled into optimized C++ code and the HMC sampler is run to generate thousands of posterior samples. This results in a full distribution of plausible parameter values, rather than just a point estimate, allowing users to compute credible intervals, visualize uncertainty, and make probabilistic predictions.

### S2 Order of acquisition diffusion analysis (OADA) likelihood

If a user only has information about the order in which events occurred, they may use an OADA model, which has a slightly different likelihood to TADA models. For each event, OADA models the probability of each individual $i$ becoming the next individual to experience an event. OADA models do not estimate an intrinsic rate, only a social transmission rate. Thus, it makes no assumptions about the shape of $\lambda_0$, but assumes that it is the same for all individuals. For more detail regarding OADA models, we direct readers to Hasenjager, Leadbeater, and Hoppitt [5] (Section 4) and Hoppitt, Boogert, and Laland [1].

We treat the likelihood of each event $e$ occurring to individual $i$ at time $t_e$ as a multinomial draw from the vector of relative hazards at that time step

$$\mathscr{L}_e = \frac{\lambda_{it_e}}{\sum_{j\in\mathcal{N}_{z_{nt_e}=0}} \lambda_{jt_e}}, \tag{10}$$

where $j$ are individuals from the full set of naive individuals at time $t_e$. The full log-likelihood of OADA models is given by

$$\log\mathscr{L}(\theta|D) = \sum_k \sum_e \left[ \log\lambda_{kit_e} - \log\left( \sum_{j\in\mathcal{N}_{z_{nt}=0}} \lambda_{kjt_e} \right) \right], \tag{11}$$

summing over trials $k$ and events $e$.

### S3 Definition of priors

STbayes departs from the previous Bayesian NBDA implementation in that priors are not entirely uninformative ($U(-10,10)$ for the intrinsic rate $log(\lambda_0)$ and social transmission rate $\log(s')$). This extreme range of values generates highly unlikely predictions in a prior predictive check (Figure S1A), and is generally not recommended for parameters fit on a log scale [6]. For example, large values of $\lambda_0$ would result in diffusions finishing within the first timestep. The following weakly informative priors produce a more plausible range of predictions for most data.

$$\log(\lambda_0) \sim N(-4,2), \tag{12}$$
$$\log(s') \sim N(-4,2), \tag{13}$$
$$B_x \sim N(0,1), \tag{14}$$
$$\Gamma_x \sim N(0,1), \tag{15}$$
$$\Phi_x \sim N(0,1), \tag{16}$$
$$\log(f) \sim N(0,1), \tag{17}$$
$$k_{\text{raw}} \sim N(0,3), \tag{18}$$
$$z_{id} \sim N(0,1), \tag{19}$$
$$\sigma_{id} \sim \text{half}N(0,1), \tag{20}$$
$$\rho_{id} \sim \text{lkj corr cholesky}(3) \tag{21}$$
$$z_{trial} \sim N(0,1), \tag{22}$$
$$\sigma_{trial} \sim \text{half}N(0,1), \tag{23}$$
$$\rho_{trial} \sim \text{lkj corr cholesky}(3) \tag{24}$$
$$\gamma \sim N(0,1) \tag{25}$$

Users may supply alternative priors as a list using the `priors` argument of `generate_STb_model()`, or saving and directly editing the STAN program before providing its file path to the `model` argument `fit_STb()`. Any priors not included in the argument will be set to default values. Prior predictive checks for standard and complex transmission are shown in Figure S1.

We set a more conservative prior on $log(f)$ than $k_{\text{raw}}$ for numerical stability. $z$, $\sigma$ and $\rho$ are fit when estimating varying effects of individuals or trials. $\rho_{id} \sim$ lkj corr cholesky($\eta$) describes the prior assumption about the correlation structure among parameters in a hierarchical model. $\eta < 1$ indicates a belief parameters are correlated, $\eta = 1$ indicates uniform prior over all possible correlations, and $\eta > 1$ indicates a belief that parameters are not correlated. $\gamma$ is defined when users fit a Weibull shaped intrinsic rate.

For convenience, here is the list of default priors providing the strings to use for each parameter:

```
default_priors <- list(
    log_lambda0 = "normal(-4, 2)",
    log_sprime = "normal(-4, 2)",
    beta_ILV = "normal(0,1)",
    log_f = "normal(0,1)",
    k_raw = "normal(0,3)",
    z_veff = "normal(0,1)",
    sigma_veff = "normal(0,1)",
    rho_veff = "lkj_corr_cholesky(3)",
    gamma = "normal(0,1)"
)
```

### S4 Guidance on MCMC model fitting

By default, the STbayes package will fit models using 1 chains of 1000 iterations (500 warm-up iterations and 500 sampling iterations per chain). Using the function `STb_summary()` on the fit will show the number of effective samples (ESS bulk, ESS tail as calculated by the posterior package) for each parameter. ESS bulk is the number of effective samples after halving the number of iterations in each chain and multiplying the number of chains by 2. ESS tail is the number of effective samples present in the 5% and 95% quantiles of the posterior, and aid users in deciding if more iterations are needed to reliably estimate tail values. More details may be found in Vehtari et al. [7].

For initial fitting of a model, the default settings should be adequate to get an impression of the posterior. Importantly, fitting one chain (rather than more than one) gives some useful diagnostic output if the chains fail (for example, specific lines in the STAN code that have caused the failure). For publication, we recommend increasing the number of chains to 4 or more. We also suggest that users run these chains in parallel if they are using a multi-core processor by using the `parallel_chains` argument.

The number of effective samples needed depends on what the users want to do (see section 9.5 of McElreath [6]). Several hundreds of ESS are enough to reliably estimate the central tendencies (mean, median) of the posterior, and reliably recovering variance parameters and/or the tails of distributions needs an order of magnitude more. Stan documentation recommends using a threshold of $ESS_{bulk} \geq 100 * n_{chains}$ as an indicator of convergence. Our summary function also includes Rhat as a convergence estimate, and users should use a threshold of $Rhat \leq 1.01$ for reliable estimates. More details for interpreting Rhat can be found in Vehtari et al. [7].

#### S4.1 Convergence failure

Model convergence indicates that the posterior distribution has been adequately sampled. A model that has not converged will not produce reliable estimates of the posterior, and it is important that users not ignore warnings from the sampler, or the diagnostics we provide in the summary function. Here, we give general guidance and preliminary steps to mitigate the problem. We strongly recommend visiting `https://mc-stan.org/learn-stan/diagnostics-warnings.html` for more detailed guidance, as well as referring to Gelman et al. [8] for best practices.

1. **Divergent transitions**: Divergent transitions happen when the posterior has extremely steep surfaces that are difficult for to explore. If there are a handful of divergent transitions, yet Rhat and ESS are otherwise fine, these may be reasonably ignored. Narrowing the priors based on domain knowledge may help resolve divergent transitions that arise from identifiability, as narrow priors are more informative and keep the sampler from straying into sharper parts of the posterior. If there are ILVs, the user can try using a normal(0,.75) or normal(0,.5) prior. Another option, although avoid this if possible, would be to increase the `adapt\_delta`, which increases the resolution at which the sampler explores. This will result in a longer time to fit the model.

2. **Rhat warnings**: Rhat compares estimates between and within chains to evaluate how well mixed the chains are. Users should aim for $Rhat \leq 1.01$ to indicate good mixing. High Rhat will usually occur along with divergent transitions, indicating a difficult to explore posterior geometry. We suggest simplifying the model as much as possible, and adding in elements to determine which variables are causing the difficult geometry. Users can also use pair-plots to identify funnel-like shapes, or multi-modal shapes in the posterior that might be causing trouble.

3. **Maximum treedepth errors**: This type of error is a sampling efficiency error, rather than a validity error. If ESS and Rhat look good, users can ignore it. Users may increase the maximum by passing the control parameter `max\_treedepth` to `STB\_fit`.

4. **BFMI low**: This means that either the warmup-phase of the chain was not sufficient, or there are some heavy tails that were not well explored. We suggest increasing the number of iterations.

We strongly suggest users read the Stan documentation by following above link before diagnosing problems. Reducing the model complexity and amount of data, and using stronger priors are all good starting points. We also invite users to simulate data using the simulation scripts provided in the reproducibility repository associated with this paper, and then fit the model to that data. Finally, please contact the corresponding author with any specific questions. Given the flexibility of the package, there may be custom cases that require directly editing the Stan code STbayes generates.

### S5 Description of %ST

We may calculate the probability of event $e$ occurring through social transmission as the social component of the rate of acquisition divided by the full rate (intrinsic and social component), at $t_e$:

$$P(\text{e occurred through ST}) = \frac{e^{\Gamma_i} s' T(a, z, w)}{\lambda_{t_e}}. \tag{26}$$

%ST is then the mean probability over all events. This can be generalized to multi-network models by calculating the probability that each event occurred through social transmission on network $n$:

$$P(\text{e occurred through ST on network n}) = \frac{e^{\Gamma_i} s' T(a_n, z, w)}{\lambda_{t_e}}. \tag{27}$$

%ST for network $n$ is again the mean probability over all events.

### S6 Description of cross validation model comparison

We fit a model to observations $y_i, ..., y_n$, representing the times at which each individual in a population experienced an event (e.g. acquiring a novel behaviour or disease). We define a set of parameters $\theta$, which include the intrinsic rate, the social transmission rate, and other covariates, and the model outputs a posterior distribution of values for each parameter $p(\theta|y)$. Leave-one-out cross validation (LOO-CV) is a technique where the model is trained on $n-1$ observations and tested on the single left-out observation, repeating this process for each data point to estimate out-of-sample predictive performance, as measured by the expected log predictive density (ELPD). This is defined as:

$$\text{ELPD} = \sum_{i=1}^{n} \log p(y_i | y_{-i}, M) \tag{28}$$

where $y_i$ represents the left-out observation, $y_{-i}$ denotes the dataset excluding $y_i$, and $M$ is the model under evaluation. The higher the ELPD, the better the model's predictive performance. However, computing ELPD directly using LOO-CV is computationally expensive, requiring the model to be refitted multiple times. WAIC estimates ELPD by incorporating both the posterior mean log-likelihood and a variance-based penalty term. LOO-PSIS provides another approximation of ELPD, but instead of using log-likelihood variance for penalization, it uses importance sampling. For a thorough description see Vehtari, Gelman, and Gabry [9].

The WAIC estimate of ELPD is defined as:

$$\widehat{\text{ELPD}}_{\text{WAIC}} = \sum_{i=1}^{n} \log\left(\frac{1}{S}\sum_{s=1}^{S} p(y_i|\theta_s)\right) - \sum_{i=1}^{n} V_{s=1}^{S}(\log p(y_i|\theta_s)) \tag{29}$$

where the first term is the posterior expectation of the likelihood and the second term is a complexity penalty representing the variance of the log-likelihood across posterior samples. This ensures that models with higher uncertainty receive poorer scores. $S$ is the number of posterior samples and $\theta_s$ represents the $s$th draw of parameter values from the posterior distribution.

The LOO-PSIS estimate of ELPD is computed as:

$$\widehat{\text{ELPD}}_{\text{LOO-PSIS}} = \sum_{i=1}^{n} \log\left(\frac{\sum_{s=1}^{S} w_i^s p(y_i|\theta_s)}{\sum_{s=1}^{S} w_i^s}\right). \tag{30}$$

In summary, it introduces a weight $w_i^s$ whose value is an approximation of the ratio of the leave-one-out posterior $p(\theta_s|y_{-i})$ and the full-data posterior $p(\theta_s|y)$. This penalizes the contributions of overly influential observations.

Both can be converted to information criterion-style scores by multiplying ELDP by -2, matching the scale of other information criteria such as AIC, such that lower values of WAIC or LOOIC indicate better predictive performance.

In order to use cross-validation methods, the log-likelihood of each observation must also be calculated for each draw $(p(y_i|\theta_s))$ and exported from the model. This is efficiently done via the generated quantities block of a Stan model, such that log-likelihoods are computed once per iteration. The boolean gq argument of the `generate_STb_model()` function allows users to specify whether or not a generated quantities block is created in the STbayes model. Since all models are fit with the same set of $n$ observations, we compare models by computing the standard error of the difference in ELPD estimates between models. For example, if one wanted to compute relative support for social transmission (equivalent to $\Delta AIC$ in the NBDA package):

$$se(\widehat{\text{ELPD}}^A - \widehat{\text{ELPD}}^B) = \sqrt{n V_{i=1}^{n}(\widehat{\text{ELPD}}_i^A - \widehat{\text{ELPD}}_i^B)}. \tag{31}$$

If the SE of a difference crosses zero, the user should be less certain that the best predictive model is any better than the alternative. This comparison can be accomplished in STbayes with `STb_compare()`, which automates the comparison between two or more models. We direct readers to the vignettes for further detail.

### S7 Description of simulations used for model validation

We validated STbayes models using two types of diffusion simulations that generated 1) more idealised data where networks were fixed, and 2) less idealised data that included motile agents.

In the first case, we defined a fixed, undirected social network prior to the start of the simulation. Each simulation trial was run on a new instantiation of a random k-regular graph of size N = 50 and degree k

= 5 (each individual has exactly 5 edges). The model proceeded in discrete time steps, with all individuals initialized as naive. At each time step, naive individuals evaluated their probability of acquiring the behaviour based on social exposure and an intrinsic rate. For each naive individual, a hazard with rate

$$\lambda_{ti} = \lambda_0 + s' T(a, z) \tag{32}$$

was calculated, where $\lambda_0$ was the intrinsic rate, and $s'$ was the strength of social influence. Transmission function $T()$ depended on whether we were validating simple or complex transmission. For simple transmission,

$$T(a, z) = \sum_j a_{tij} z_{tj}. \tag{33}$$

For complex transmission, we used either f-complex or k-complex equations. The per-step learning probability was calculated as $1 - exp(-\lambda_{ti})$, and individuals independently drew a Bernoulli trial to determine whether they learned in that time step. Once informed, individuals remained informed, and their time of acquisition was recorded. The process continued until all individuals were informed or the maximum number of time steps was reached.

In order to validate that STbayes could recover varying effects of individuals, we simulated 10, 50 and 100 diffusion trials from the same 50 individuals with known values for intrinsic and social transmission rates, and normally distributed variation by individual for each. The model recovered main and varying effects. Accuracy of varying effects estimates predictably increased with the number of trials.

#### S7.1 Motile agent simulations

We adapted the simulation model from Chimento and Farine [3] to simulate transmission in populations of dynamically moving populations. Explicitly encoding the time it takes to move from one position to the other changes transmission dynamics, and potentially can make it very difficult to infer true parameter values from static representations of networks, which is what is often done by empirical researchers. We used these simulations to test the improvement gained by including dynamic transmission weights and dynamic networks, as well as the "high-resolution" mode. We simulated transmission in populations of $N = 50$ agents, who were initialized with random starting positions on a toroidal surface of 80x80 units. If agents were within 15 units of one another as measured by euclidean distance, they were considered connected, and behavioural productions of one agent would potentially influence an acquisition event in another agent.

Each timestep, agents updated their positions according to a movement rule. We chose to use nomadic movement, where agents move largely in straight lines, but occasionally change heading. Nomadic movement creates a sparse instantaneous network, but a dense static representation of the network, likely making dynamic networks useful for inference. If agents were knowledgable, they would emit cues according to a stochastically changing probability, meaning that they would use known behaviour variably, making dynamic transmission weights useful. Naive agents that were connected were influenced by the emissions of knowledgable agents, and could potentially acquire the behaviour through social transmission. We tested a positive frequency-dependent rule, as well as a standard simple transmission rule.

The simulations ran until all agents were knowledgable. We recorded the times at which each agent experienced the event. For networks, we recorded an overall static representation of the network over the course of the simulation, a dynamic network that summarized connections between inter-event intervals, and a high-resolution network that recorded connections at each timestep. We recorded the same types of data for transmission weights. Using this data, we could then fit successively complicated models presented in Section 7.

#### S7.2 Parameter values

For all simulations, we set set $log(\lambda_0) = -6$ and $log(s') = -4$. For the validation of random effects, we set $\sigma_{log(\lambda_0)} = 0.25$, $\sigma_{log(s')} = 0.25$. For complex transmission tests, $k = -0.75$ for conformity, $k = 0.75$ for anti-conformity.

#### S7.3 Sample sizes

Stochastic simulation variance can dominate the estimates of NBDA type models for single trials. Furthermore, the number of subjects, as well as network density can significantly impact model power. For static network simulations, we standardized both. We found that a significant number of trials were needed to estimate varying effects of individuals (Figure S7 versus Figure 3), testing $N = 10$, 50 and 100 trials for the varying effects model presented in Section 3. We used $N = 10$ trials for the simulations testing edge uncertainty presented in Section 5. We used $N = 10$ trials for the simulations of complex transmission on static networks for the models presented in Section 6. We used $N = 10$ trials for the simulations using motile agents presented in Section 7.

### S8 Power analysis

There are two important power tests for researchers to consider. The first is the number of observations needed to correctly support a model that includes social transmission, versus an asocial only null model. The second is the number of observations needed to correctly infer the strength of social transmission relative to the intrinsic rate ($s$). The number of individuals present in the population, as well as the network configuration, both affect the relative power of STbayes to determine either. To help users make an informed decision about sample size, we perform power analyses using different combinations of population size ($N \in \{12, 24, 36, 48, 60\}$ with different numbers of trials $(1, 5, 10, 15, 20)$. On a third dimension, we test different network densities: dense networks ($k = N/3$), medium-dense networks ($k = N/4$) and sparse networks ($K = N/6$). For each combination, we simulated 10 diffusions using a small effect of social transmission ($s = 2$).

To check whether models correctly support social transmission, we fit both an asocial only model and a full model, and calculated the relative support using $\Delta ELPD_{loo}$. We then checked whether the median upper 95% CI of $\Delta ELPD_{loo}$ for the asocial model crossed 0. If it is below zero, this indicates that the researcher could conclude that social transmission was supported. We find that few trials are needed to correctly support social transmission under a small effect of $s = 2$. 1 trial was sufficient for population sizes greater than or equal to 24 (Figure S5. We do recommend performing multiple trials when possible, and maximizing the number of subjects in each trial when possible. Figure S4 illustrates the runtime under different combinations of population size and number of trials. Network density will not affect runtime as these calculations are vectorized. For performing your own power analysis, we direct you to the vignette on the topic in the online documentation.

### S9 Description of frequency dependent transmission equations

The transmission function of "f" transmission is

$$T(a_i, z, w, t) = \frac{\sum_j a_{ij} z_{jt} w_{jt}{}^f}{\sum_j a_{ij} z_{jt} w_{jt}{}^f + \sum_j a_{ij}(1 - z_{jt})^f}. \tag{34}$$

The frequency-dependent bias parameter $f$ is constrained as $f \geq 0$.

The transmission function of "k" transmission is based on another tuneable sigmoid function that translates the proportion of "knowledgable" or "infected" associates into a normalized output $y$

$$x = \frac{\sum_j a_{ij} z_{jt} w_{jt}}{\sum_j a_{ij} z_{jt} w_{jt} + \sum_j a_{ij}(1 - z_{jt})}$$
$$x' = 2x - 1$$
$$T(a_i, z, w, t) = \frac{\left(\frac{x' - kx'}{k - 2k|x'| + 1} + 1\right)}{2}.$$

The transformation depends on parameter $k$, which is constrained between $(-1, 1)$.

### S10 Supplementary Figures

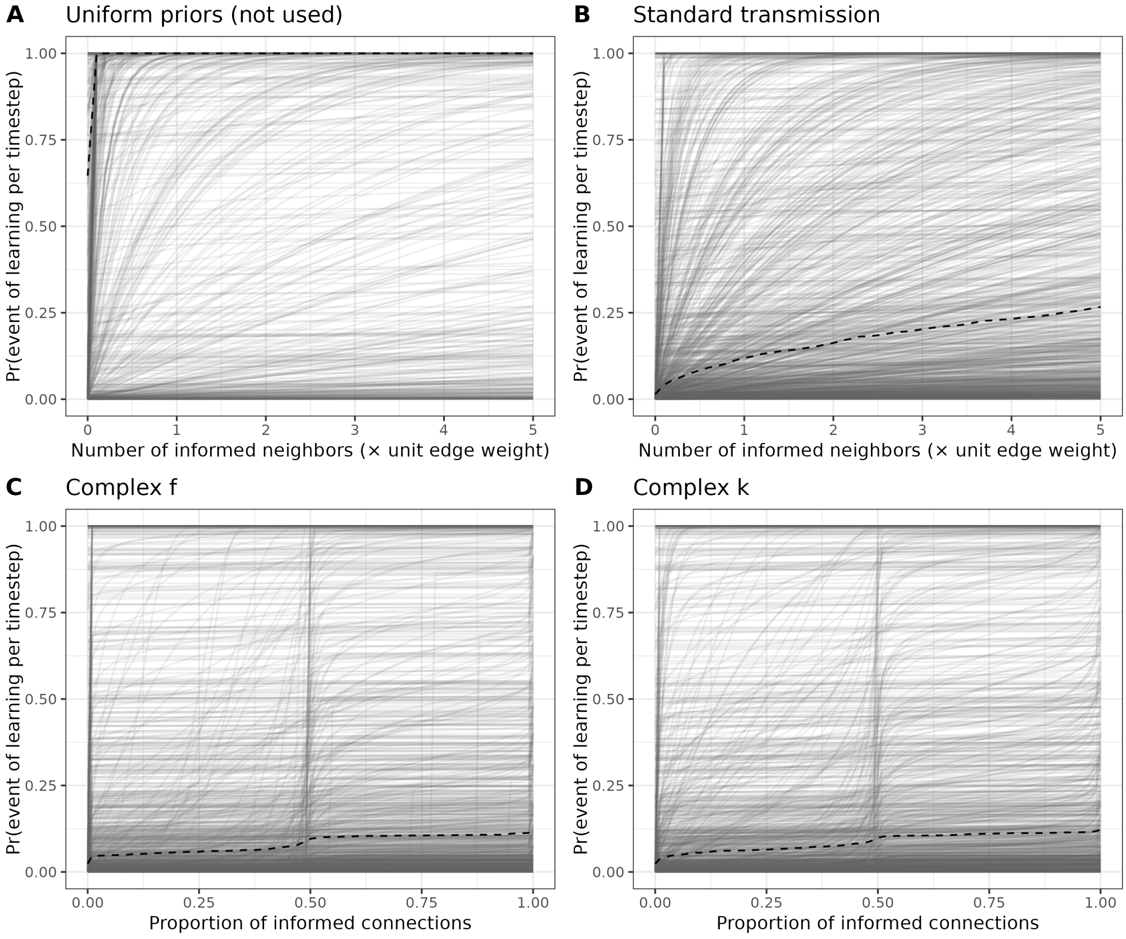


Figure S1: Prior predictive check. Predicted relationships under the default priors of STbayes between the number of informed connections (x-axis) and the instantaneous probability of an event happening (y-axis). Median values are shown with dashed lines. A) Suggested uniform priors from the previous implementation of Bayesian NBDA produce a large number of unlikely relationships. Our weakly informative priors provide a more credible relationship for B) standard transmission, C) complex f transmission and D) complex k transmission.

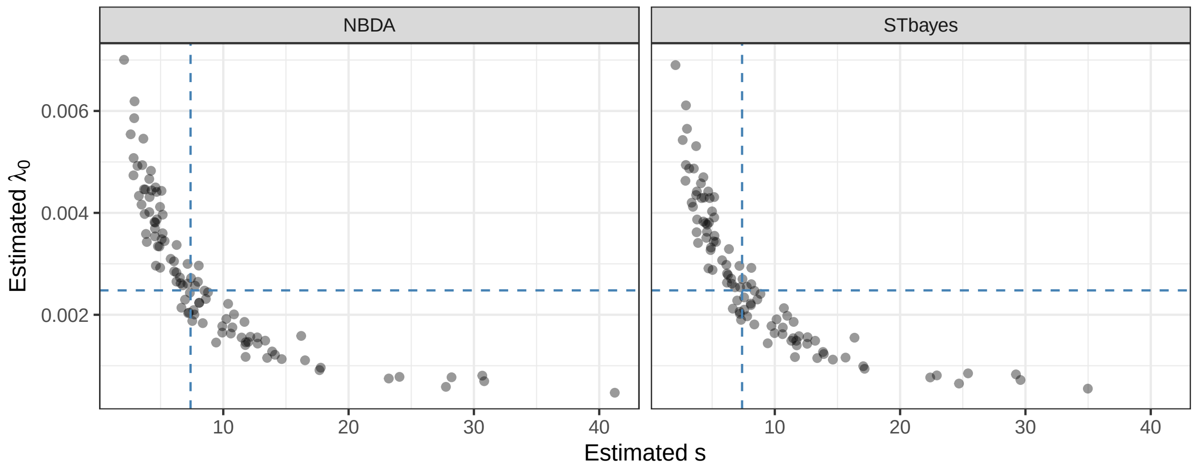


Figure S2: Comparison of estimated parameters between NBDA and STbayes. Shown are the point estimates for either $s$ (x-axis) or $\lambda_0$ (y-axis) using either package (panel) from single simulations. Blue dashed lines indicate the true values of parameters used for all simulations. Both packages produce a similar distribution of estimates.

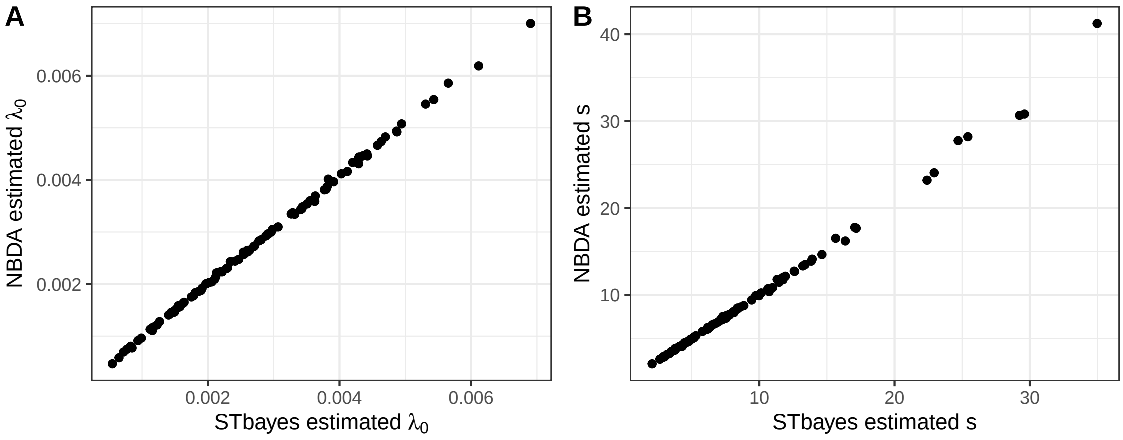


Figure S3: Correlation between NBDA and STbayes parameter estimates. For each simulated dataset where $\lambda_0 = 0.001$ and $s = 5$, we compare the estimated values from STbayes (x-axis) and NBDA (y-axis).

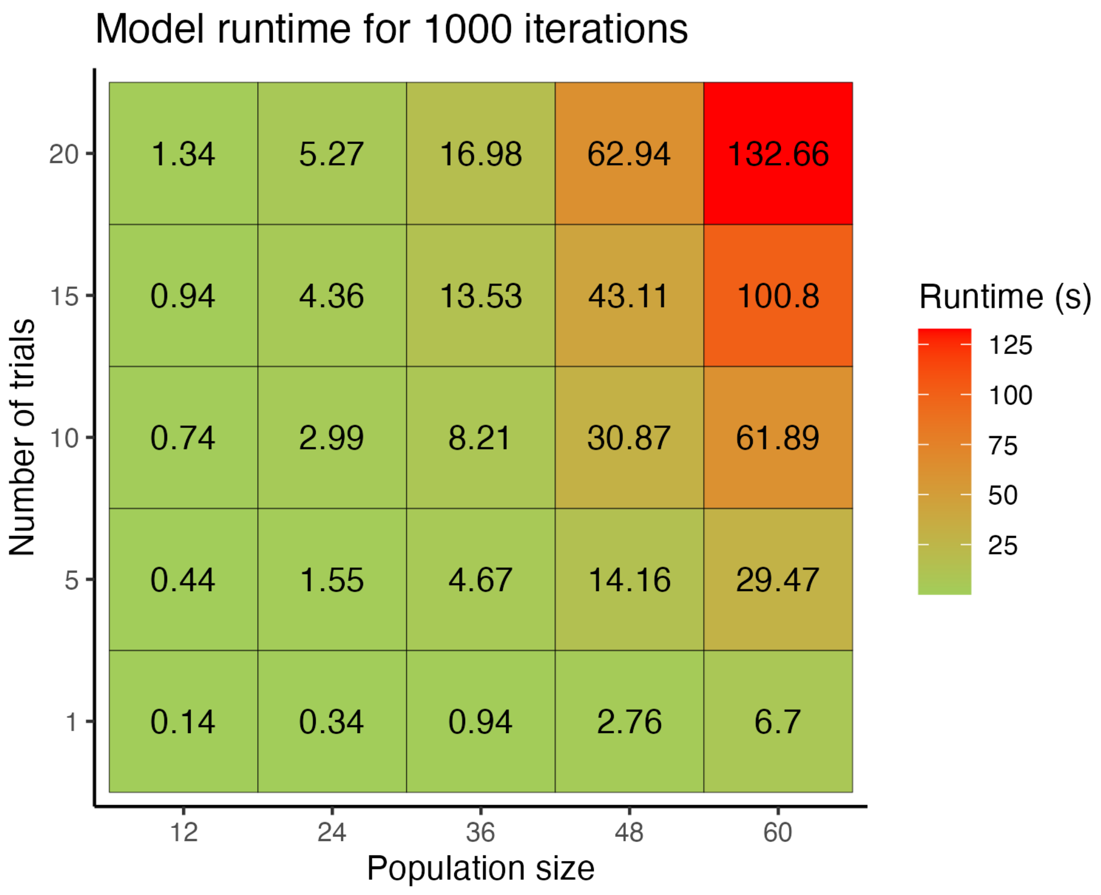


Figure S4: Model runtimes. A heat-map illustrating the time taken to fit models with varying population size (x-axis) and number of trials (y-axis) in seconds. All models were fit with the standard transmission function, no varying effects, and no ILVs. Including more parameters will increase the runtime. All models were fit with default settings of 1 chain and 1000 iterations. The number of chains should be increased to 4 or more for final model runs, and we encourage users to run these in parallel using the parallel_chains argument.

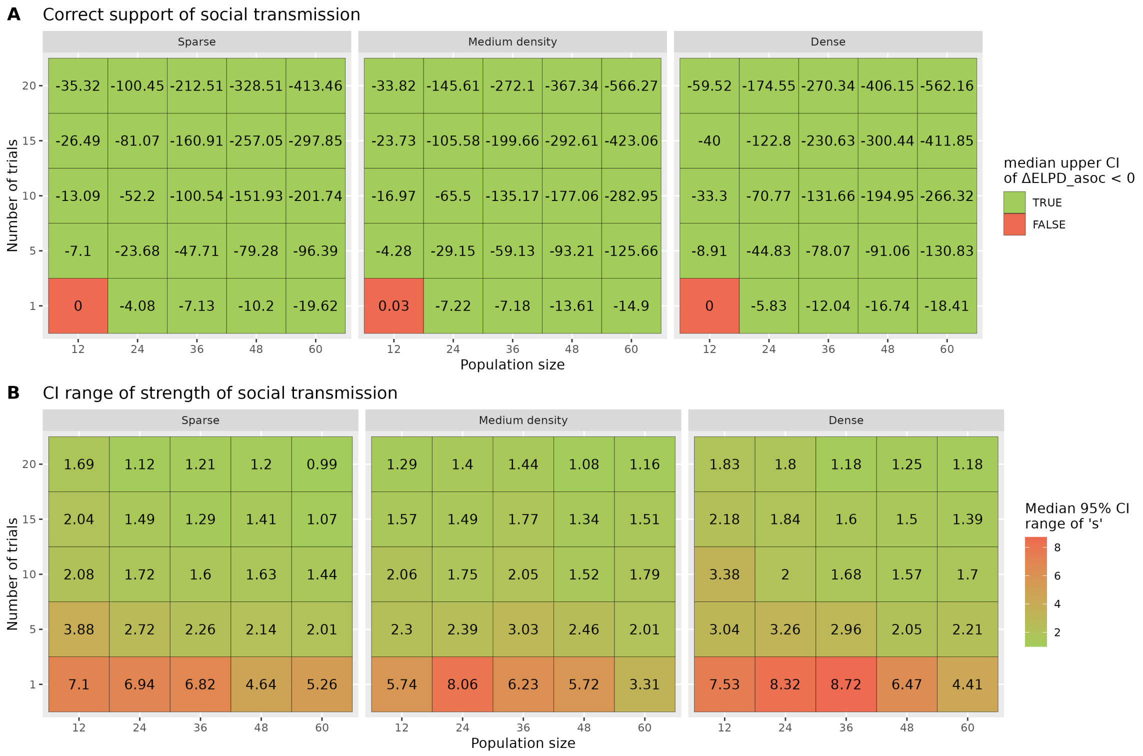


Figure S5: Power analysis results. Presented are metrics from networks of 3 different densities, with different population sizes and number of trials. Density is scaled to the population size. A) Few trials are needed to reliably support social transmission over asocial only models. B) The 95% CI of estimated $s$ decreases with population size and number of trials. Researchers who need a more confident estimate of $s$ should design experiments with multiple trials.

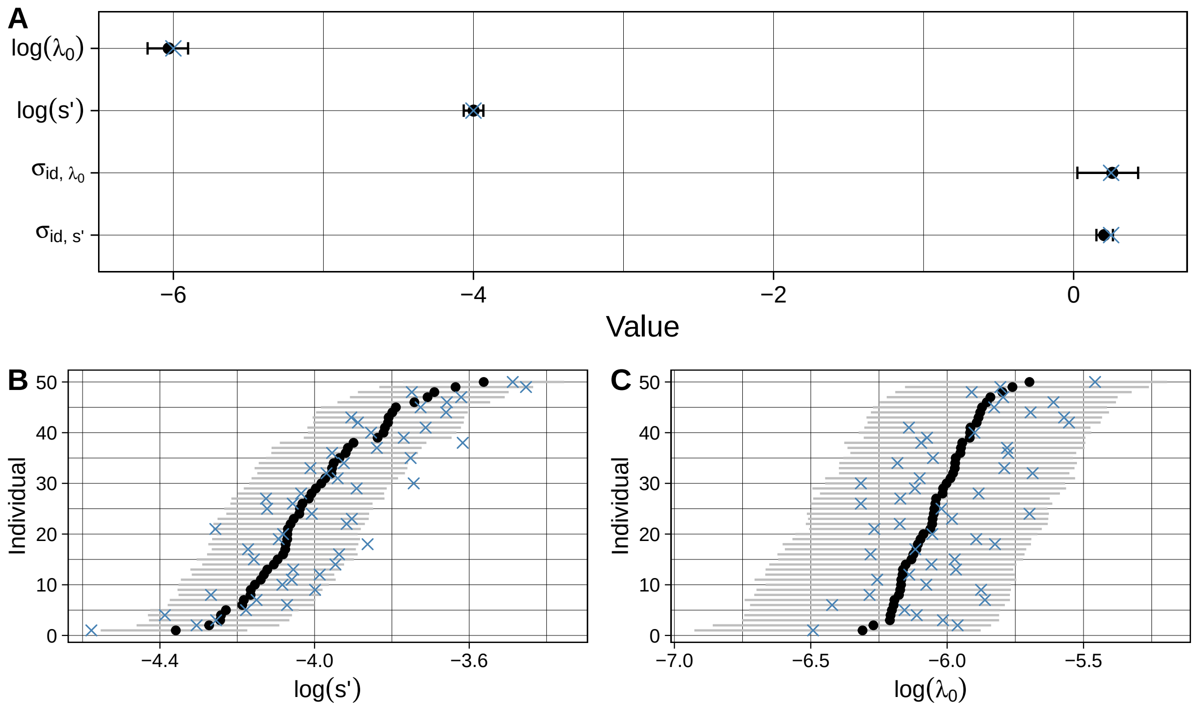


Figure S6: STbayes estimates of main and varying effects. We simulated 100 diffusion trials from the same 50 individuals with varying values of intrinsic rate and strength of social learning. A) Main effect estimates, black error bar and point are 95% HPDI and median, blue x is the true value. Varying effects of individuals for B) log $s'$ and C) log $\lambda_0$, comparing the true value (x) with the estimated value and 95% CI.

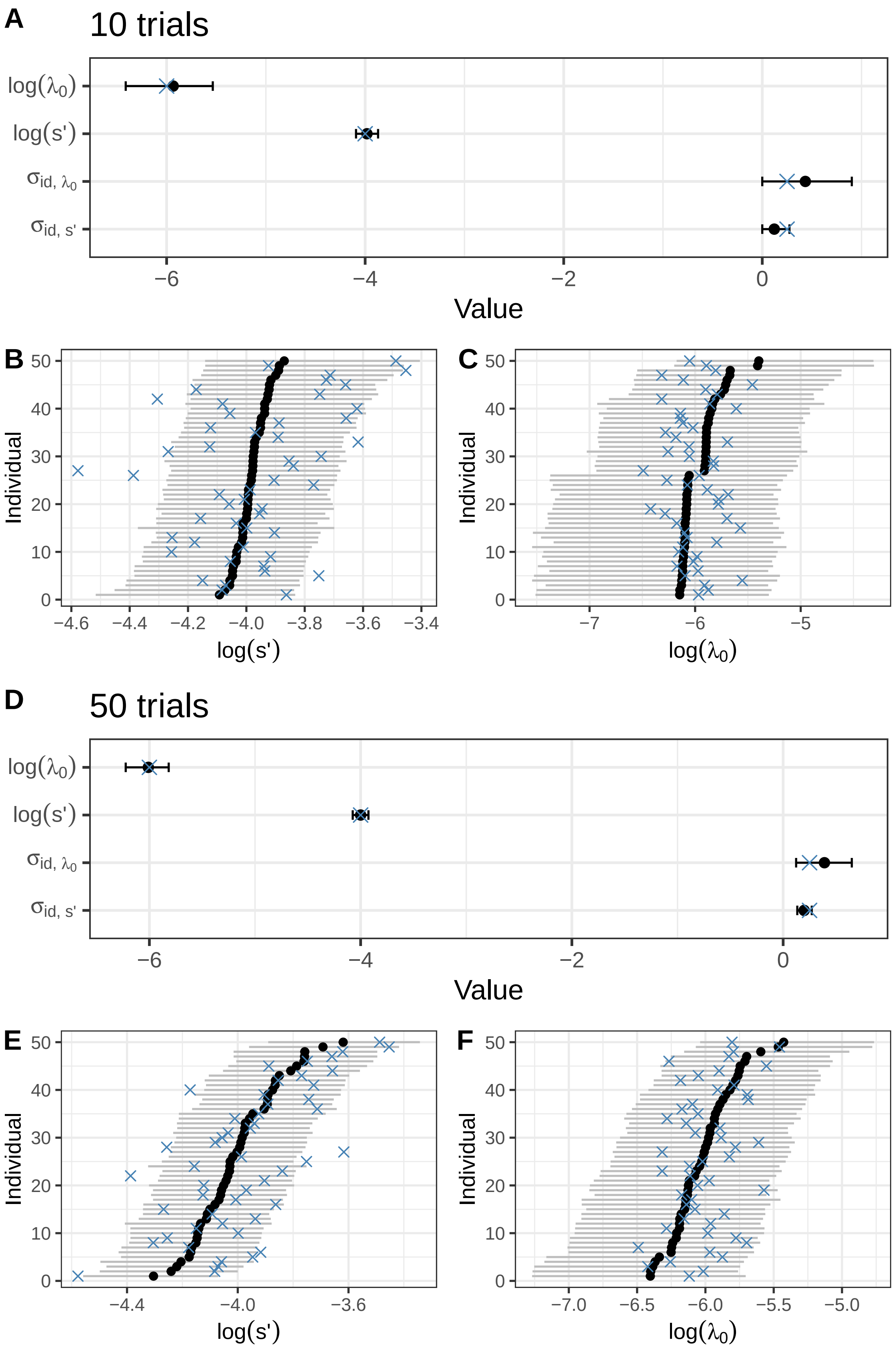


Figure S7: Varying effects estimates with a lower sample size. We additionally fit a model with varying effects for $\lambda_0$ and $s'$ on a subset of 10 (panels **A, B, C**) and 50 trials (panels **D, E, F**). True values are shown with blue "x"s, with median and 95% HPDI as black point-ranges in all panels. 10 trials was sufficient to recover main effects, although a number of true varying effects fell outside of the CI.

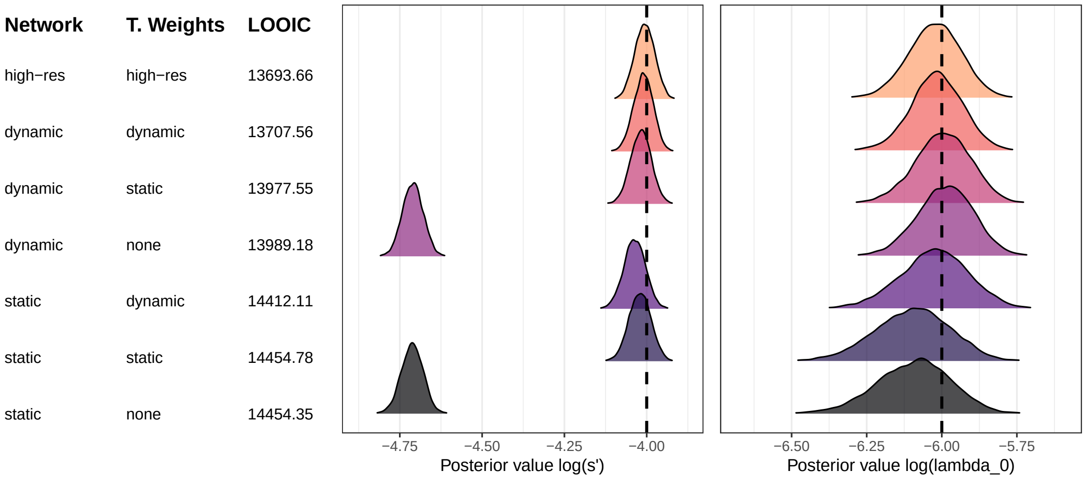


Figure S8: **Testing dynamic transmission weights and networks under simple transmission.** We simulated diffusions using a standard "simple" transmission rule on dynamic networks of motile agents. Compared to complex transmission presented in the main text, parameters underlying simple transmission processes are easier to recover without ultra-high-resolution data.

#### S11 Supplementary Tables

| Notation | Name | Description |
|---|---|---|
| $\lambda_i$ | hazard rate | the rate at which individual $i$ experiences the target event (e.g. acquisition of behavior, pathogen, information). |
| $\lambda_0$ | intrinsic rate (also called baseline rate [5]) | the underlying rate of a target event in the absence of social transmission. |
| $s'$ | social transmission rate | the rate of transmission per unit connection. |
| $s$ | strength of social transmission | the rate of transmission per unit connection relative to the intrinsic rate. |
| $z_i$ | event state | describes the state of individual $i$ ($0 =$ event has not occurred, $1 =$ event has occurred). |
| $a_{ij}$ | edge weight | describes the edge between individuals $i$ and $j$ within a network. |
| $B_i$ | intrinsic rate ILV | linear combination of individual-level variables affecting the intrinsic rate. |
| $\Gamma_i$ | transmission rate ILV | linear combination of individual-level variables affecting the social transmission rate. |
| $\Psi_i$ | multiplicative ILV | linear combination of individual-level variables affecting both the intrinsic and social transmission rate. |
| $w_j$ | transmission weight | describes the rate of a cue of interest (e.g. expression of target behaviour) is exhibited by individual $j$. |

Table S1: Summary of notation.

| Parameter | Median | MAD | HPDI_Lower | HPDI_Upper | n_eff | Rhat |
|---|---|---|---|---|---|---|
| log_lambda_0_mean | -6.031 | 0.068 | -6.172 | -5.902 | 8102.993 | 1.000 |
| log_s_prime_mean | -3.999 | 0.033 | -4.065 | -3.934 | 3704.247 | 1.001 |
| lambda_0_mean | 0.002 | 0.000 | 0.002 | 0.003 | 8103.006 | 1.000 |
| sprime_mean | 0.018 | 0.001 | 0.017 | 0.020 | 3704.233 | 1.001 |
| s_mean | 7.632 | 0.598 | 6.510 | 8.876 | 6840.528 | 1.001 |
| sigma_ID[1] | 0.258 | 0.097 | 0.025 | 0.430 | 2950.694 | 1.001 |
| sigma_ID[2] | 0.204 | 0.028 | 0.152 | 0.262 | 4131.197 | 1.001 |
| percent_ST[1] | 0.874 | 0.003 | 0.868 | 0.880 | 16510.609 | 1.000 |

Table S2: Typical summary statistics from STb_summary() function. Output taken from the varying effects model shown in Figure S6. Columns include parameter names, median value and median average divergence (MAD), upper and lower limits of the credible interval (defaults to 95% HPDI), number of effective samples (n_eff), and Rhat diagnostics. Log and linear values of $\lambda_0$ and $s'$ are always given. The strength of social transmission ($s = s'/\lambda_0$) and %ST are calculated in the generated quantities block. Parameters sigma_ID[1] and sigma_ID[2] are the SD of varying effects for $\lambda_0$ and $s'$. For multi-network models, the summary function will show multiple $s$ and %ST values associated with each network. The order will be identical to the columns in the networks dataframe.

| Parameter | Median | MAD | CI_Lower | CI_Upper | ess_bulk | ess_tail | $\hat{R}$ |
|---|---|---|---|---|---|---|---|
| log_lambda_0_mean | -5.903 | 0.061 | -6.026 | -5.789 | 4636.0 | 5282.5 | 1.000 |
| log_s_prime_mean | -3.999 | 0.025 | -4.047 | -3.946 | 4750.6 | 5151.3 | 1.001 |
| lambda_0 | 0.003 | 0.000 | 0.002 | 0.003 | 4636.0 | 5282.5 | 1.000 |
| s | 6.715 | 0.497 | 5.751 | 7.681 | 4014.8 | 4395.8 | 1.000 |
| s_prime | 0.018 | 0.000 | 0.017 | 0.019 | 4750.4 | 5151.3 | 1.001 |
| percent_ST[1] | 0.781 | 0.007 | 0.767 | 0.795 | 4014.8 | 4395.8 | 1.000 |

Table S3: Summary of posterior estimates for model where edges are point estimates (from section 5).

| Parameter | Median | MAD | CI_Lower | CI_Upper | ess_bulk | ess_tail | $\hat{R}$ |
|---|---|---|---|---|---|---|---|
| log_lambda_0_mean | -5.975 | 0.062 | -6.103 | -5.856 | 16304.3 | 5541.1 | 1.000 |
| log_s_prime_mean | -3.953 | 0.035 | -4.022 | -3.883 | 7668.2 | 6341.7 | 1.000 |
| lambda_0 | 0.003 | 0.000 | 0.002 | 0.003 | 16304.2 | 5541.1 | 1.000 |
| s | 7.565 | 0.578 | 6.453 | 8.759 | 11889.4 | 5701.9 | 1.001 |
| s_prime | 0.019 | 0.001 | 0.018 | 0.021 | 7668.2 | 6341.7 | 1.000 |
| percent_ST[1] | 0.797 | 0.007 | 0.783 | 0.810 | 13580.1 | 5271.3 | 1.001 |

Table S4: Summary of posterior estimates for model where edge uncertainty is included (Section 5).

| Parameter | Median | MAD | CI_Lower | CI_Upper | ess_bulk | ess_tail | $\hat{R}$ |
|---|---|---|---|---|---|---|---|
| log_lambda_0_mean | -5.990 | 0.102 | -6.197 | -5.800 | 6722.3 | 7399.6 | 1.001 |
| log_s_prime_mean | -4.099 | 0.072 | -4.246 | -3.960 | 6915.3 | 7669.7 | 1.001 |
| k_raw | -2.470 | 0.344 | -3.221 | -1.806 | 7320.8 | 6123.8 | 1.000 |
| lambda_0 | 0.003 | 0.000 | 0.002 | 0.003 | 6722.3 | 7399.6 | 1.001 |
| s_prime | 0.017 | 0.001 | 0.014 | 0.019 | 6915.4 | 7669.7 | 1.001 |
| k_shape | -0.844 | 0.049 | -0.939 | -0.746 | 7320.8 | 6123.8 | 1.000 |
| s | 6.632 | 0.981 | 4.879 | 8.707 | 5981.6 | 6918.7 | 1.001 |
| percent_ST[1] | 0.578 | 0.028 | 0.522 | 0.630 | 5556.9 | 6149.7 | 1.001 |

Table S5: Summary of posterior estimates for complex transmission using a static network (Section 6).
